# Bivariate multilevel meta-analysis of log response ratio and standardized mean difference for robust and reproducible environmental and biological sciences

**DOI:** 10.1101/2024.05.13.594019

**Authors:** Yefeng Yang, Coralie Williams, Alistair M. Senior, Kyle Morrison, Lorenzo Ricolfi, Jinming Pan, Malgorzata Lagisz, Shinichi Nakagawa

**Affiliations:** Evolution & Ecology Research Centre and School of Biological, Earth and Environmental Sciences, University of New South Wales, Sydney, NSW 2052, Australia; Charles Perkins Centre and School of Life and Environmental Sciences, The University of Sydney, Sydney, NSW 2006, Australia; Department of Biosystems Engineering, Zhejiang University, Hangzhou 310058, China

**Author notes:** Correspondence Shinichi Nakagawa, Evolution & Ecology Research Centre and School of Biological, Earth and Environmental Sciences, University of New South Wales, Sydney, NSW 2052, Australia, Jinming Pan, Department of Biosystems Engineering, Zhejiang University, Hangzhou 310058, China. Senior author.

**Keywords:** research synthesis, effect size, meta-analysis, meta-regression, random-effects model, mixed-effects model, hierarchical linear model, multivariate analysis, selective reporting, p-hacking

## Abstract

Meta-analytic modelling plays a pivotal role in synthesizing research and informing relevant policies. Yet researchers face many analytical challenges. In environmental and biological sciences, one of the most common yet unrecognised issues is the selection between two common effect size metrics, log response ratio (lnRR) and standardized mean difference (SMD); these two are the most popular and alternative effect sizes. Having to choose between them creates room for analytical flexibility, which is susceptible to researcher degrees of freedom. Another common issue is failure to deal with statistical dependence between effect sizes, resulting in invalid inferences on evidence. We propose addressing these two issues through the joint synthesis (dual use) of lnRR and SMD. Using 75 meta-analyses, including 3,887 environmental/biological primary studies (∼20,000 effect sizes), we show a high false positive rate (40%) in conventional meta-analytic practices (random-effects model) compared to the proposed bivariate multilevel meta-analysis of lnRR and SMD along with robust variance estimation. Relying solely on either lnRR or SMD results in non-trivial discrepancies in detecting statistically significant effects (18%) and occasional inconsistencies in sign (9%). Discrepancies in interpreting effect size, heterogeneity, and publication bias are prevalent between models using lnRR and SMD (e.g., 52% for publication bias). In contrast, bivariate synthesis of lnRR and SMD yields substantial information gain, reducing standard error in effect size estimates by 29%, equivalent to adding 40 additional effect sizes. We present a user-friendly website with a step-by-step implementation guide. Our proposed robust approach aspires to improve meta-analytic modelling using lnRR and SMD in environmental and biological evidence synthesis, amplifying their reproducibility and credibility.

## 1 INTRODUCTION

Meta-analytic modelling represents an indispensable methodology within the evidence synthesis ecosystem, transforming evidence-based research and practice across diverse fields (Gurevitch et al., 2018). In global change biology, meta-analytic modelling serves three key purposes: (1) drawing generalizations about the overall patterns of environmental and biological phenomena (i.e., the overall effect), (2) assessing the degree of (in)consistency of phenomena across contexts (i.e., heterogeneity), and (3) identifying informative predictors of phenomena (i.e., moderators). Methodological innovation yields diverse statistical techniques that have led to influential meta-analytic discoveries, which have shaped scientific investigation and policy (Jackson et al., 2011; Nakagawa et al., 2012; Ogle et al., 2021; Riley et al., 2017). Nonetheless, the breadth of statistical techniques available creates room for analytic flexibility or “researcher degrees of freedom” (Parker et al., 2023; Steegen et al., 2016). This flexibility in the choice of analytical approach raises the possibility that analysts may (un)intentionally search methodological specifications to align with their desired findings, leading to confirmation bias (i.e., seeing what they want to see) and the risk of erroneously concluding the null effect as the true effect (i.e., a high false positive rate). Notably, this concern extends to the most basic methodological decision: the selection of effect size metrics.

The selection of effect size metrics is inherently linked to the underlying hypothesis being tested in a meta-analysis. For example, when examining the hypothesis that “artificial light at night suppresses the melatonin secretion” (Yang et al., 2024), effect size metrics based on mean differences, rather than correlations, become a logical selection. This is because these metrics intuitively articulate the hypothesis by expressing the difference in mean outcomes between treatment (exposure) and control (no exposure) groups. Among the effect size metrics available for comparing the means of two groups, log response ratio (lnRR; Hedges et al., 1999) and standardized mean difference (SMD; Hedges, 1981), emerge as the two most employed metrics (Nakagawa et al., 2023).

However, a less recognized aspect is that lnRR and SMD are based on different assumptions regarding the environmental and biological processes, potentially leading to different conclusions. Specifically, lnRR assumes a multiplicative effect of the treatment on the control group’s mean outcomes, measuring the percentage change in mean outcomes in the treatment group compared to the control group (Figure 1).

**Figure 1.**
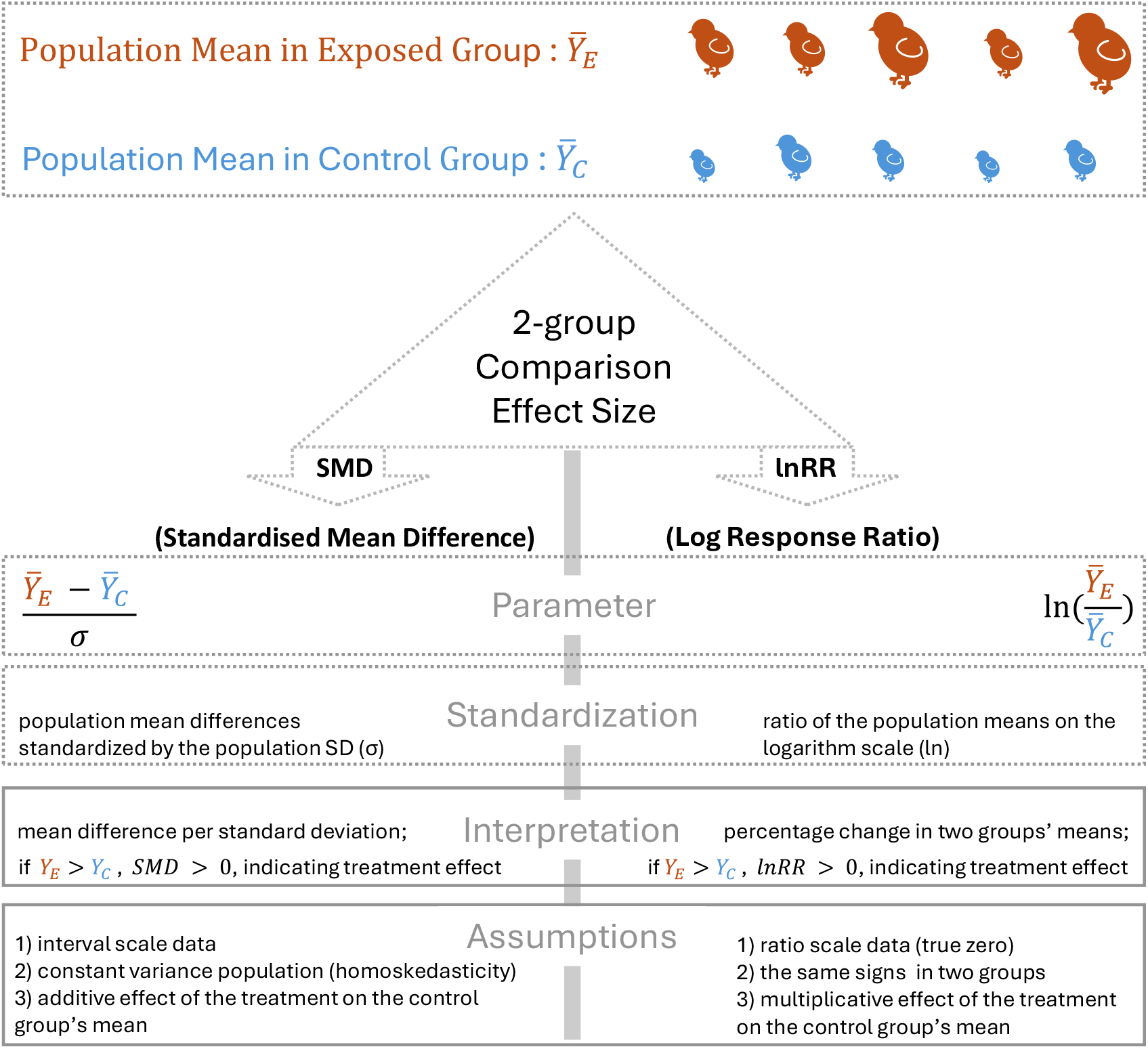
Characteristics of the two most commonly usesd effect size measures used for comparing the means of two groups: log response ratio (lnRR) and standardized mean difference (SMD).

Conversely, SMD assumes an additive effect of the treatment on the control group’s mean outcomes, quantifying mean differences between the treatment and control groups. While SMD standardizes mean differences against the population standard deviation, this standardization imposes challenges in interpreting the observed mean difference as the causal strength of the environmental and biological process. This is because the population standard deviation may co-vary with experimental design factors (Osenberg et al., 1997). For example, field studies have consistently demonstrated larger variances compared to laboratory studies (Hillebrand et al., 2014). Consequently, when the conclusions derived from these two effect size metrics are contradictory, analysts may choose the metric favouring the hypothesis being tested.

Another important methodological consideration involves overlooking the statistical independence among effect sizes, a critical assumption for valid statistical inferences (Borenstein et al., 2010). In environmental and biological sciences, fixed-effect (FE) and random-effects (RE) models are frequently employed (Nakagawa et al., 2023). However, a meta-analysis often includes studies reporting multiple effect sizes, and thus each effect size does not contribute unique information (a scenario also known as effect size dependence or multiplicity). Indeed, a survey of meta-analyses in environmental sciences revealed that 100% of them involved multiple effect sizes per study (Nakagawa et al., 2023). Consequently, the assumption of statistical independence is violated by ‘conventional’ FE and RE models, resulting in underestimated uncertainty, artefactually narrow confidence intervals, inflated p values, and incorrect inferences regarding general patterns of an effect of interest (Van den Noortgate et al., 2013). Fortunately, various strategies have been developed to address statistical dependence in effect sizes. A single effect size per study can be generated by aggregating or selecting one effect size estimate to satisfy the assumption of statistical independence (*ad hoc* strategies; Cheung, 2014). More sophisticated techniques, such as multilevel and multivariate modelling and robust variance estimation, directly account for the statistical dependence of multiple effect size estimates from a single study but also offer additional insights (Jackson et al., 2011; Ogle et al., 2021; Van den Noortgate et al., 2013; Yang, Macleod, et al., 2023). Despite the elegance and information gain provided by these advanced techniques, their uptake has been relatively low and slow in practice (Nakagawa et al., 2023).

To address the above challenges, we propose the joint synthesis (dual use) of lnRR and SMD. Our objectives are as follows: (1) assessing how often researcher degrees of freedom, in terms of choosing meta-analytic models, may have contributed to the rejection of null hypotheses, revealing potentially erroneous conclusions, (2) investigating the extent of disparities between meta-analyses utilizing lnRR and SMD, and (3) exploring the additional biological and statistical insights gained from the joint meta-analysis of lnRR and SMD using our approach. The following section describes the distinct statistical and biological properties of lnRR and SMD. Building upon the conventional meta-analytic techniques, we introduce the theoretical underpinnings of our method. Subsequently, using 75 meta-analyses from environmental and biological sciences, we demonstrate the necessity and advantages of our approach in two aspects. Specifically, we first empirically examine the discrepancies between lnRR and SMD in terms of the overall effect estimation (e.g., inconsistency in magnitude and sign), heterogeneity quantification, and publication bias test. Secondly, we quantitatively measure the biological and statistical information gain resulting from the application of our method. We also provide a website showing the step-by-step implementation of our proposed methods using readily available software (e.g., R package metafor; Viechtbauer, 2010): https://yefeng0920.github.io/BRMA_tutorial_git/. Our approaches aim to reduce researcher degrees of freedom in making methodological decisions for conducting meta-analyses, thereby increasing the robustness and reproducibility of resultant evidence synthesis.

## 2 MATERIALS AND METHODS

### 2.1 Mean difference effect size metric

In the context of meta-analysis, the term “effect size” carries three interpretations (Nakagawa et al., 2007), referring to either (1) a parameter, (2) an estimator, or (3) an estimate. The effect size parameter denotes a numerical metric, index, or measure that quantifies the direction and magnitude of the treatment effect resulting from a global change driver, such as temperature increase or acidification. The effect size estimator represents the estimation of the treatment effect based on sample estimates (e.g., means and standard deviations) of population parameters from either laboratory or field study data. The effect size estimate is a value, indicating the empirical direction and magnitude of the treatment effect. While we use the three definitions interchangeably throughout the paper, it is necessary to clarify them in the context of two commonly used mean difference effect size metrics: log response ratio (lnRR; Hedges et al., 1999) and standardized mean difference (SMD; Hedges, 1981).

#### 2.1.1 Log response ratio (lnRR)

The lnRR is an effect size metric (Hedges et al., 1999), quantifying the strength of an environmental and biological intervention by comparing the population means of a focal outcome between a group exposed to an environmental stressor (e.g., a global change driver or treatment in general) and a control group (benign environment). The mathematical definition of the lnRR effect size parameter is given by Equation 1 (the first definition; Hedges et al., 1999):

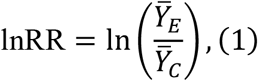

where In () denotes the natural logarithm function, 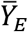 and 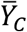 represent the mean level of the focal outcome from an environmental/biological stressor group (subscript *E*) and control group (subscript *C*), respectively. In practical scenarios, the population means (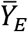 and 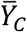 ) are seldom known and must be estimated from sample data obtained in laboratory or field studies. A simple way to circumvent the unknown population means is to plug their sample means (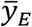 and 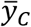 ) derived from the empirical studies. This results in a basic lnRR effect size estimator:

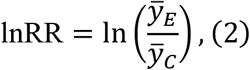

Assuming the focal outcome measurements are statistically independent, the estimator of sampling (error) variance of the lnRR can be obtained by the delta method:

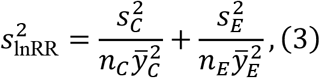

where *s*_*C*_ and *s*_*E*_ denote the sample standard deviations corresponding to sample means (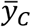 and 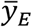), *n*_*C*_ and *n*_*E*_ are the respective sample sizes (the number of replicates).

In cases where a study indicates no environmental or biological effect (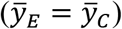), the lnRR value is zero, assuming a large sample size to mitigate statistical noise. For smaller sample sizes, bias-corrected estimators for the lnRR and 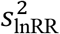, have been proposed (Lajeunesse, 2015; Pustejovsky, 2018). We note that the assumption of independent sampling error is often violated (Nakagawa et al., 2023), impacting the accuracy of the sampling variance estimation and potentially inflating Type I error rates. We will introduce a method named robust variance estimation (RVE) to account for unknown dependence in sampling error which we present later (Hedges et al., 2010). The lnRR effect size parameter is widely favoured due to its superior biological and statistical properties. One notable biological property is its conversion into a proportional change metric: 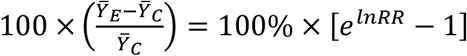 (where *e* is the exponential function), providing an intuitive interpretation of the strength environmental or biological effect in terms of percentage change in the focal outcome measurements induced by an environmental stressor. However, a limitation of the lnRR is that it can only be applied to ratio scale measurements (positive mean values) because of the properties of the natural logarithm function, ln (the natural logarithm of zero and negative numbers is undefined). Additional properties will be discussed in the context of empirical comparisons with the SMD effect size.

#### 2.1.2 Standardized mean difference (SMD)

The SMD is an effect size that quantifies the strength of the environmental effect by comparing the population means of a focal outcome between a group exposed to an environmental stressor group and a control group, standardized by the population standard deviation of the focal trait. The mathematical definition of the SMD effect size parameter is given by Equation 4 (Hedges, 1981):

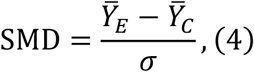

where *σ* denotes the population standard deviation of the focal trait, assumed to be constant across groups (*σ* = *σ*_*E*_ = *σ*_*C*_), and other notations are the same as above. As with lnRR, since population parameters (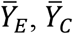, and *σ*) are generally unknown, we need an estimator of the SMD effect size parameter:

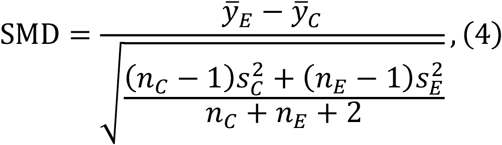

where all notations are the same as above. Equation 4 is known as Cohen’s *d*, where the denominator is called the pooled standard deviation *s*_*pooled*_, calculated by weighting the sample deviations of the two groups by their degrees of freedom (Hedges, 1981). The pooled standard deviation *s*_*pooled*_ serves as a proxy for *σ*. Assuming statistically independent sampling errors, the estimator of the sampling (error) variance of the SMD is given by the delta method:

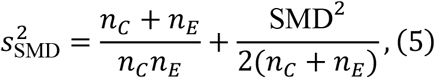

If a laboratory or field study yields an estimate of SMD of one, it implies that the mean level of the focal outcome in the environmental stressor group is one sample standard deviation away from that in the benign environment group (or control group). A small-sample-size bias-corrected estimator for the SMD and 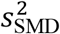 is also available and is known as Hedge’s g (Hedges, 1981). The SMD effect size parameter possesses several advantageous biological and statistical properties, such as expressing the strength of the environmental/biological pressure in the unit of standard deviation.

This feature allows the inclusion of primary studies using different types of traits and measurement instruments. We note that some properties of SMD often lead to discrepancies with lnRR in terms of meta-analytic results, a topic we delve into in

## RESULTS AND DISCUSSION

### 2.2 Conventional meta-analysis analysis comparing the mean of two groups

The fixed-effect (FE) and random-effects (RE) models represent conventional approaches in the meta-analysis (Borenstein et al., 2010), and are employed in over 70% of studies in environmental sciences, including global change biology (Nakagawa et al., 2023). Consider *J* studies being included for meta-analysis, each *j*-th study contributing only one effect size estimate with corresponding sampling variance.

Typically, either lnRR or SMD serves as the effect size metric. The RE model, employing one of these metrics, is given by:

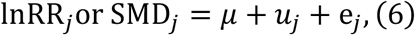

where the InRR_*j*_ (or SMD_*j*_) denotes the effect size estimate in *j*-th study, *μ* represents the average effect size parameter (also known as the overall effect), *u*_*j*_ is the studyspecific random deviation of the true effect (i.e., parameter) in the *j*-th study from *μ*, and e_*j*_ denotes the sampling error describing the difference between the effect size estimate (Equation 2) and the effect size parameter for *j*-th study (Equation 1). The standard statistical assumptions of Equation 6 are: *u*_*j*_*∼N*(0, *τ*^2^), and 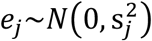, where *N*() represents the normal distribution. When *τ*^2^ = 0, *u*_*j*_ = 0 for all *J* studies, the RE model (Equation 6) simplifies to the FE model, assuming homogeneity of true effects. However, these conventional approaches can suffer from the issues outlined in **INTRODUCTION** when effect sizes are correlated and/or either lnRR or SMD are used for analysis.

### 2.3 Multilevel meta-analysis comparing means of two groups

The first approach involves using the three-level multilevel meta-analytic model (MLMA) to synthesize lnRR along with SMD. The basic MLMA of lnRR and SMD is the three-level hierarchical models that describe the effect size estimates, InRR_*ij*_ and SMD_*ij*_, at the sampling level, within-study level, and between-study level (Van den Noortgate et al., 2013). Consider a meta-analysis with *J* studies, each *j*-th study contributing *n*_*j*_ effect size estimates and sampling variances. The mathematical representation of the first approach is given by:

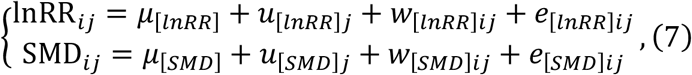

where *μ* denotes the overall effect (with the subscript distinguishing the lnRR and SMD), *u* denotes a between-study level random error (random-effect) term, *w* denotes a within-study level random error term, *e* is the corresponding sampling error. All three error terms are typically assumed to follow the normal distributions to ensure the unbiased estimation of the average effect size parameter: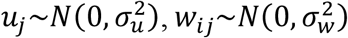, and 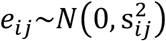, where 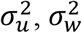, and 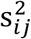 denote the between-study variance, within-study variance, and sampling variance.

The first approach based on Equation 7 possesses two properties that make it an attractive strategy for environmental and biological meta-analysis. Statistically, the nested random effect structure accounts for the correlation between the true effect sizes (see below), ensuring valid statistical inference of the overall effect (Van den Noortgate et al., 2013). Biologically, the total heterogeneity can be partitioned into between- and within-study level heterogeneity (measured as *I*^2^; see technical details in **Supplementary Materials**), allowing for the identification of moderator variables corresponding to different contextual characteristics. More importantly, between- and within-study-level heterogeneity carries important biological information. For example, a small magnitude of between-study level heterogeneity means that the effect sizes are consistent at the study level, indicating that the study characteristics do not systematically affect the magnitude of the effect sizes, and thus the corresponding research findings are generalizable across different studies. In contrast, a large magnitude of with-study level heterogeneity means that the true effect sizes vary within the same study, and with-study characteristics can be used to explain the heterogeneity.

Under the framework of MLMA, the variance-covariance matrix of the true effect sizes (InRR_*ij*_ or SMD_*ij*_) is expressed as:

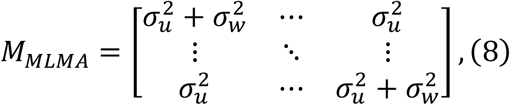

The *M*_*MLMA*_ can be reparametrized as a compound symmetric random-effects structure within the framework of the multivariate meta-analysis (MVMA; see **Section 2.4**):

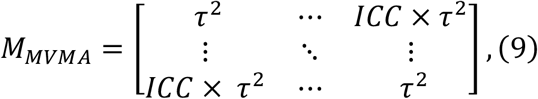

where 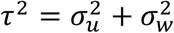 represents the variance in the true effect, and 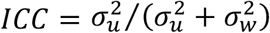 denotes the intraclass correlation coefficient, indicating the strength of dependence/correlation between the true effects within the same study (details see **Supplementary Materials**). Although the MLMA assumes independent sampling errors 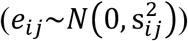, it can account for the correlated sampling errors by incorporating a sampling variance-covariance matrix. Such a sampling variance-covariance matrix has a similar structure as that in the MVMA. Assume a metaanalysis involves two studies where the first study contributes two effect size estimates due to shared control and the second study contributes one effect size estimate. Then, the sampling variance-covariance matrix has the following form:

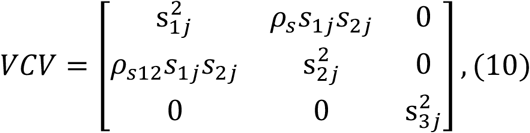

To relax the independence assumption of the true effect size and sampling error, we will introduce how to use the robust variance estimation (RVE) technique to relax it in **Section 2.5**.

### 2.4 Bivariate multilevel meta-analysis comparing means of two groups

The second advanced approach we propose is the use of a bivariate multilevel metaanalytic model (BMLMA) to simultaneously synthesize the lnRR and SMD. The BMLMA, or more broadly, a multivariate multilevel meta-analytic model, is a bivariate meta-analytic model with a multilevel random-effects structure. It can jointly model multiple types of outcomes and multiple sources of heterogeneity (Jackson et al., 2011), simultaneously accounting for statistical dependence due to both the multilevel and correlated nature of effect sizes. Although it is rarely employed in meta-analytic practices within environmental and biological sciences due to conceptual complexity and implementation difficulties, its potential merits are substantial (Yang, Macleod, et al., 2023). To have an intuitive understanding of the BMLMA, imagine extending conventional meta-analytic models to bivariate versions, analogous to extending analysis of variance (ANOVA) to multivariate analysis of variance (MANOVA).

Mathematically, the second approach is expressed as:

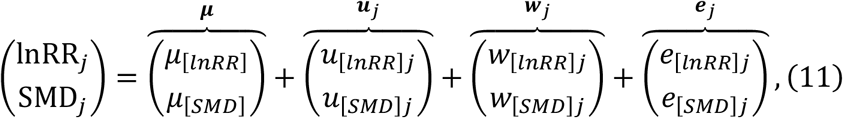

where ***μ*** is a bivariate of the overall effect representing the overall effect corresponding to lnRR and SMD, ***u***_*j*_ is a 2 *×* 1 vector of between-study level random effects for the lnRR and SMD, *w*_*j*_ is a 2 *×* 1 vector of within-study level random effects for the lnRR and SMD, and *e*_*j*_ is a 2 *×* 1 vector of sampling errors. Under typical statistical assumptions, ***u***_*j*_, *w*_*j*_, and *e*_*j*_ follow the multivariate normal distributions: ***u***_*j*_*∼MVN*(0, ***T***^2^), *w*_*j*_*∼MVN*(0, ***Γ***^2^), and 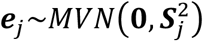. The betweenand within-study variance parameter ***T***^2^ and ***Γ***^2^ have the following variance-covariance structures that need estimation:

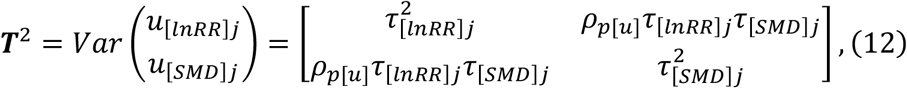

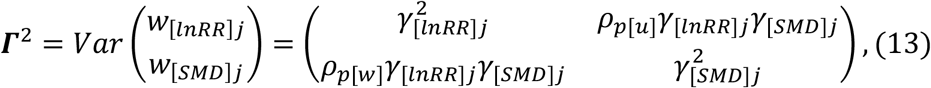

The sampling variance-covariance parameter 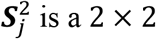 is a 2 *×* 2 symmetric matrix of the *j*th study, already known from the corresponding estimator (Equations 3 and 5):

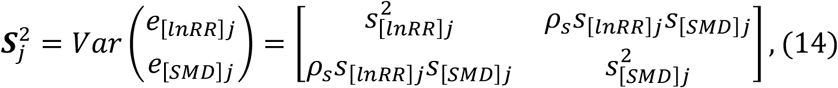

The correlated random-effect structures (Equations 12 and 13) and sampling error (Equation 14) can account for the statistical dependence in the true effect size and sampling error, resulting in more accurate statistical inference on the overall effect. The proposed approach exhibits two useful properties for environmental and biological meta-analysis. First, the utilization of the correlation *ρ*_*p*_ between the lnRR and SMD parameters can increase the precision of the overall effect estimation, a concept known as borrowing of strength (BoS; Copas et al., 2018; Riley et al., 2017). This is particularly valuable for the joint synthesis of lnRR and SMD when the effect size estimator of lnRR or SMD becomes invalid for some included studies, such as when sample mean values for environmental/biological interventions and control groups have opposite signs or one of the means is zero, leading to missing effect sizes. Second, a low estimated correlation between lnRR and SMD parameters indicates that a large true effect quantified using lnRR (or SMD) is not necessarily meant to imply a large true effect quantified using SMD (or lnRR). In such cases, results corresponding to both effect size metrics should be transparently reported and carefully interpreted.

Nonetheless, the proposed BMLMA approach has three limitations. Firstly, the sampling correlation *ρ*_*s*_ is rarely reported in practice, often requiring the use of an educated guess or arbitrary value (e.g., 0.5 or 0.8 Pustejovsky et al., 2022; Yang, Macleod, et al., 2023). We can the RVE approach as a solution to remedy this issue later (see **Section 2.5**). Secondly, there is no established method for quantifying heterogeneity (variability around the true effect size) in the context of multivariate meta-analysis. Therefore, the heterogeneity estimates provided by the proposed BMLMA approach need to be interpreted with caution. In contrast, the first approach, MLMA of lnRR along with SMD can provide accurate heterogeneity estimates (see **Supplementary Materials**). Thirdly, the BMLMA approach is highly parameterized, and the model parameters might not be identifiable (i.e. resulting in convergence issues) when the number of studies is small.

### 2.5 Robust variance estimation (RVE)

As mentioned, although the two proposed approaches for the dual synthesis of lnRR and SMD exhibit promise; however, they have limitations under specific circumstances. One notable limitation arises from the potential underestimation of the standard error (*SE*(*μ*)) of the overall effect (*μ*) when sampling correlation (*ρ*_*s*_) is either ignored or assigned an arbitrary value (see above). Such circumstances can lead to inflated Type I error rates and potentially result in misleading conclusions. In such a case, it is necessary to complement the two proposed approaches with robust variance estimation (RVE) techniques (Pustejovsky et al., 2022; Yang, Macleod, et al., 2023). The rationale behind RVE techniques is straightforward. These techniques assume that estimates of study-specific average effects follow a distribution with an unknown (co-)variance (Hedges et al., 2010). Subsequently, RVE calculates the weighted sample standard error as an empirical estimate of the *SE*(*μ*). The mathematical formula of RVE can be found in **Supplementary Materials**. Therefore, the RVE-based standard error *SE*(*μ*) is robust to an arbitrary sampling correlation *ρ*_*s*_ and inaccurate estimates of variance components (e.g., 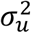, or *τ*^2^). Additionally, small sample-corrected estimators for *SE*(*μ*) are available (Joshi et al., 2022; Pustejovsky et al., 2018; Tipton, 2015). Nonetheless, we recommend examining the sensitivity of the overall effect *μ* and standard error *SE*(*μ*) to the choice of different *ρ*_*s*_.

### 2.6 Empirical performance assessment

To address the three objectives outlined in Section 1 **(INTRODUCTION)**, we compiled a database of environmental and biological meta-analyses and evaluated our proposed first and second advanced approaches using various performance metrics.

#### 2.6.1 Data source

The meta-analysis database included 75 publicly available meta-analytic datasets related to global change biology, ecology, and evolutionary biology. We collected meta-analytic datasets from Hillebrand et al. (2020), initially used to examine thresholds and regime shifts associated with various anthropogenic global change drivers, including climate warming, acidification, and exotic invasions. To enhance the comprehensiveness of our database, we also included meta-analytic datasets from ecology and evolutionary biology (Senior et al., 2016; Yang, Sánchez-Tójar, et al., 2023). The computation of lnRR and SMD for some datasets was infeasible, resulting in missing effect size estimates. For example, zero sample means could yield an estimate of infinite lnRR. After removing all missing data, our database encompassed 3,887 studies, incorporating 19,521 lnRR effect size estimates and 19,824 SMD effect size estimates. Each study contributed approximately six effect sizes, and there was a moderate correlation between them (intra-class correlation 0.45 for lnRR and 0.69 for SMD; see **Supplementary Materials**), underscoring the prevalent statistical dependence within our database.

#### 2.6.2 Prevalence of erroneous meta-analytic evidence

Extensive simulations and empirical comparison have shown that the multivariate and multilevel meta-analytic models have nominal Type I error rates in the presence of statistical dependence (Cheung, 2013; Kalaian et al., 1996; Van den Noortgate et al., 2013). In contrast, conventional meta-analytic models (by default ignoring statistical dependence) could not control Type I error rates, and thus inflate two-sided p-value. We empirically quantified the conditions wherein our proposed method indicates no overall effect *μ*_[lnRR]_ (as evidenced by a two-sided p-value < 0.05), while conventional meta-analytic models (e.g., RE model) find a statistically significant effect *μ*_[lnRR]_ (the proportion of falsely rejecting the null hypothesis *H*_0_: *μ*_[lnRR]_ = 0). We used the *rma ()* function from the *metafor* package (Viechtbauer, 2010) to fit the RE model for each of the 75 datasets, using restricted maximum likelihood (REML) as the variance estimator. For the MLMA and BMLMA, we used *rma*.*mv()* function with REML as the variance estimator. We set the quasi-Newton method as the optimizer to maximize the likelihood function over variance estimation, with a threshold of 10^-8^, a step length of 1, and a maximum iteration limit of 1000.

#### 2.6.3 Discrepancies in meta-analytic results between the log response ratio (lnRR) and standardized mean difference (SMD)

We employed three metrics related to inferential errors to evaluate the discrepancy between lnRR and SMD in terms of key meta-analytic results, including the overall effect, heterogeneity quantification (expressed as *I*^2^) and publication bias (detected by the multilevel version of Egger’s test; details see **Supplementary Materials**).

Detectability discrepancy was used to assess whether lnRR and SMD result in different pieces of statistical evidence when lnRR and SMD are jointly modelled using our proposed method. For example, in a given meta-analysis, lnRR may detect a statistically significant overall effect, while SMD might not, or *vice versa*. The sign inconsistency was used to evaluate whether lnRR and SMD yield different signs for the overall effect. The magnitude discrepancy was used to gauge whether lnRR and SMD yield different magnitudes for the overall effect or heterogeneity. Differences in magnitudes could lead to distinct interpretations, especially when compared against effect size and heterogeneity benchmarks (Correll et al., 2020; Higgins et al., 2003).

#### 2.6.4 Statistical and biological information gain

As mentioned above, the MLMA and BMLMA approaches offer potential statistical and biological information gain. To quantify the statistical information gain, we employed the BoS statistic (Copas et al., 2018). The BoS statistic measures the reduction in uncertainty, or incremental precision, of the overall effect estimate resulting from the joint synthesis of lnRR and SMD. It is calculated as 100(1 − *E*)%, where *E* represents efficiency and is given by 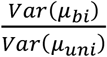 (with the numerator and denominator denoting the variance of the overall effect size estimate obtained from bivariate and univariate analyses of lnRR and SMD, respectively). We calculated BoS using the BMLMA of lnRR and SMD. To provide a more intuitive interpretation, we converted the BoS to the number of additional effect sizes augmented due to the reduction in uncertainty: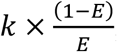, where *k* is the number of effect sizes included in each dataset (Riley et al., 2017). For quantifying biological information gain, we decomposed heterogeneity (measured as *I*^2^; see technical details in **Supplementary Materials**) at different levels using the MLMA of lnRR together with SMD.

## 3 RESULTS AND DISCUSSION

### 3.1 High prevalence of erroneous meta-analytic evidence in conventional metaanalytic approach

Our results revealed that the conventional approach (random-effects, RE, model) had consistently lower p values of the overall effect compared to the two proposed approaches, when statistical dependence was present. Given that simulations and empirical comparison have shown that the RE model inflates Type I error rates in the presence of statistical dependence, the RE model is prone to incorrectly concluding a null effect as a statistically significant overall effect (*μ*_[lnRR]_ ≠ 0 or *μ*_[*SMD*]_ ≠ 0).

Specifically, within the 31 meta-analyses where the multilevel meta-analytic (MLMA) model alongside robust variance estimation (RVE) identified a statistically nonsignificant overall effect for lnRR (*μ*_[lnRR]_ = 0), the RE model yielded 13 (42%) cases with a statistically significant overall effect (Figure 2). Similar results were observed for the MLMA model using SMD (46%; Figure 2). The bivariate multilevel metaanalysis (BMLMA) model plus RVE nullified more than 40% of initially statistically significant effects for both lnRR and SMD (Figure 2).

**Figure 2.**
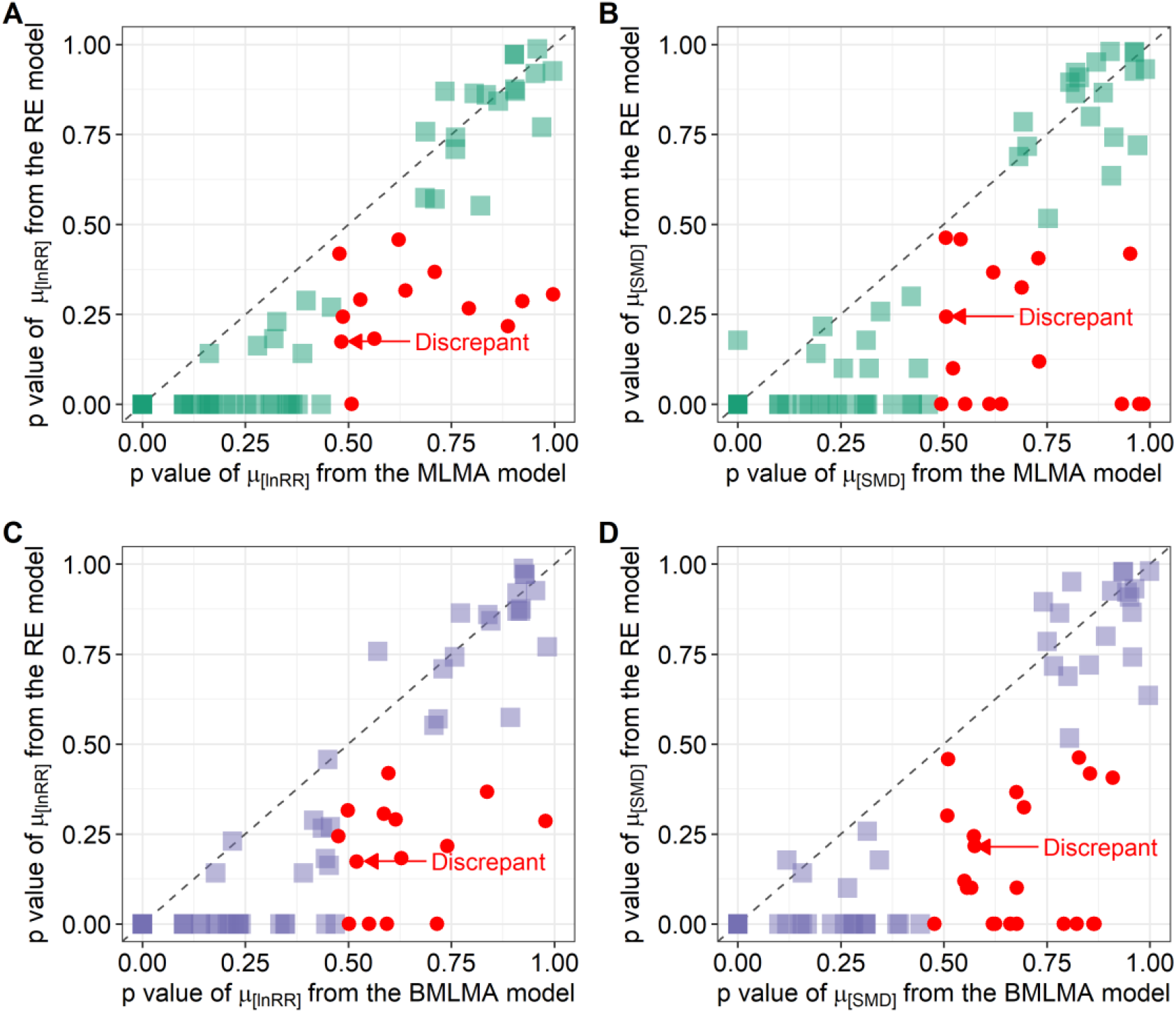
An empirical comparison of the p values of overall effect *μ* using log response ratio (lnRR) and standardized mean difference (SMD) as the effect size metric. The diagonal line signifies equality between the two metrics. The fourth root of the p value is employed for graphical representation across a broad range from 0 to 1, centering on p values between 0.05 and 0.10. The plot based on the original scale p values is reported in Figure S1. Notably, the conventional random-effects (RE) model exhibited consistently lower p values for lnRR compared to the proposed multilevel meta-analysis (MLMA; the first approach) (Panel A) and the bivariate multilevel meta-analysis (BMLMA; the second approach) (Panel C). Both proposed approaches are implemented alongside robust variance estimation (RVE) to maintain a nominal Type I error rate and ensure statistical validity of the p values. Therefore, the p values from the RE model are likely to be inflated, leading to wrong meta-analytic evidence. Similar patterns were observed in the results for SMD (Panels B and D). Red points represent the discrepancy between meta-analysis using lnRR and SMD in terms of the null hypothesis significance test of the overall effect.

These observations suggest that at least 40% of initially statistically significant overall effects became non-significant after using the MLMA or BMLMA models to account for statistical dependence. Consequently, relevant published meta-analyses using the RE model drew erroneous conclusions by incorrectly concluding the null effect as evidence of environmental and biological effects. Collectively, this implies that the conventional approach, particularly in the context of statistically dependent effect sizes, should be strongly avoided in environmental and biological meta-analysis (Nakagawa et al., 2012). Nonetheless, the subsequent findings underscore that relying solely on one effect size measure to draw inferences on the overall effect can still lead to erroneous conclusions.

### 3.2 Discrepancies between the log response ratio (lnRR) and standardized mean difference (SMD)

We found that the use of lnRR or SMD alone led to non-trivial and even large discrepancies in detectability and magnitude, and slight discrepancies in the sign of the overall effect. While the MLMA model using lnRR identified 44 cases of statistically significant overall effects, and 40 cases of statistically significant overall effects for SMD (Figure 3), only 36 cases showed statistically significant effects simultaneously, indicating a detection discrepancy of 18% (8/44) for SMD and 10% (4/40) for lnRR. Similar patterns were observed for the BMLMA using lnRR and SMD (Figure S2). In cases where SMD showed a ‘large’ overall effect (SMD = 0.2, 0.5, and 0.8 corresponding to “small”, “medium”, and “large” effects; Correll et al., 2020) based on effect size benchmarks (21 cases), lnRR ranged from 0.122 to 2 (median 0.47), corresponding to a mean percentage ranging from 11% to 339% (median 46%; Figure 4). Consequently, lnRR and SMD exhibited clear divergence in interpreting the strength of the overall effect in certain instances, as it is unreasonable to interpret 11% changes in mean difference as a large effect. Notably, we identified around 10% of cases where lnRR and SMD showed opposite signs for the overall effect derived from both the MLMA and BMLMA models (Figure S3).

**Figure 3.**
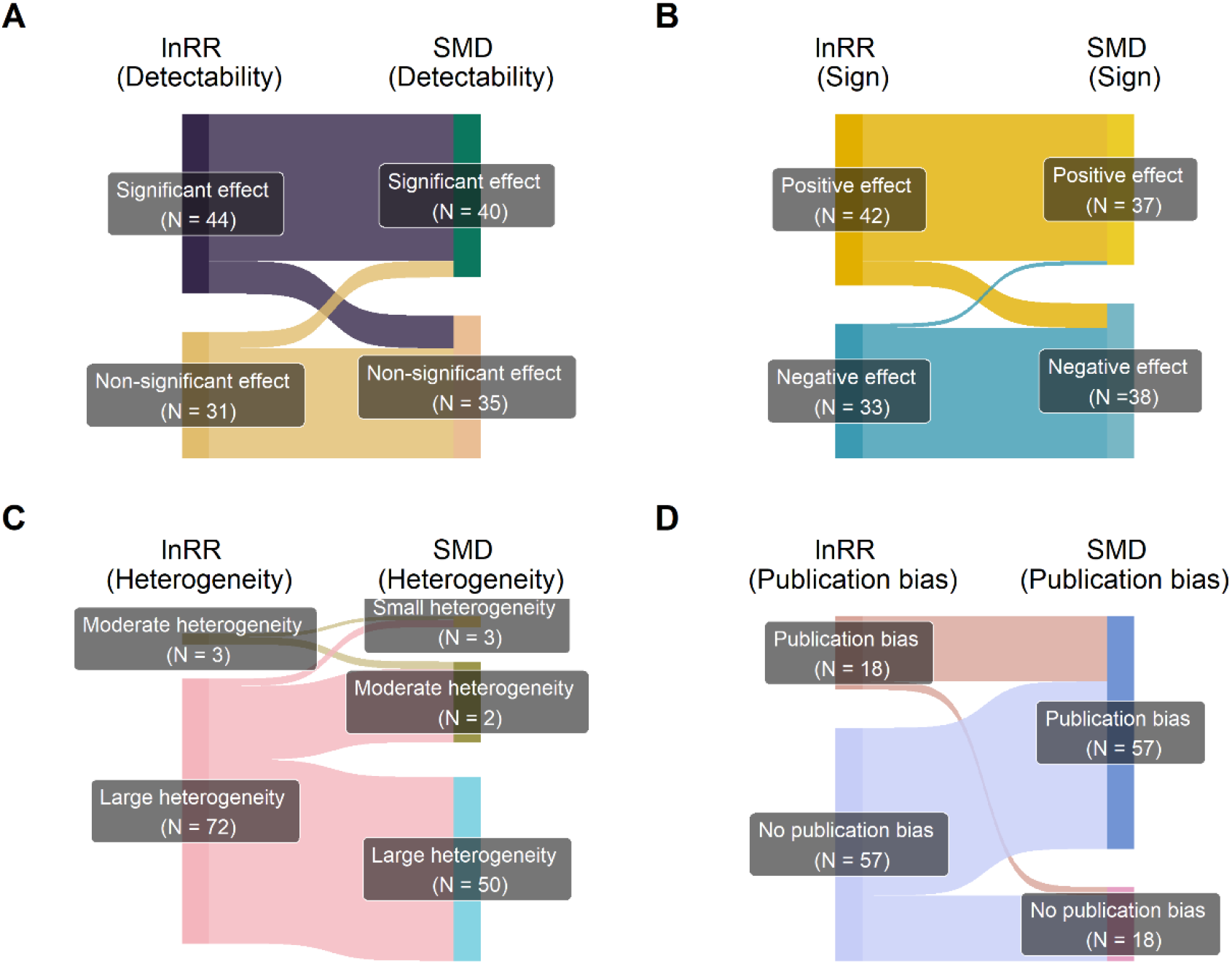
Inconsistencies in results between lnRR and SMD from the multilevel meta-analysis model (MLMA). We examined the results corresponding to the main procedures of a metaanalysis, including the null hypothesis significance test (detectability; Panel A) and sign (Panel B) of the overall effect, heterogeneity interpretation (Panel C), and publication bias test (Panel D).

**Figure 4.**
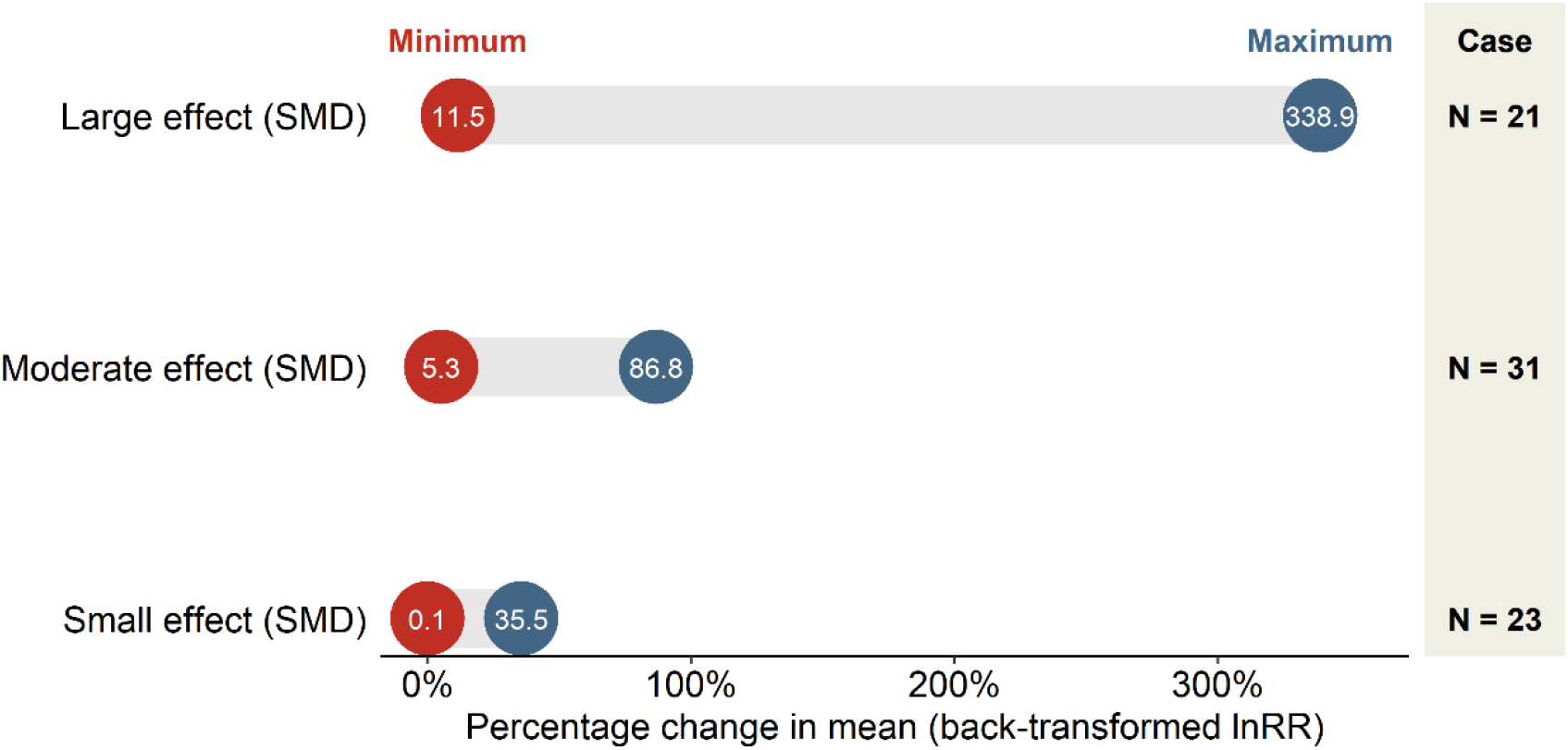
Divergent interpretations of the magnitude of the overall effect between lnRR and SMD from the multilevel meta-analysis model (MLMA). While the use of SMD suggests a large effect size, the use of lnRR implies a small effect, characterized by a modest percentage change (e.g., 11.4%) in the mean difference between the environmental or biological intervention group and the control group.

Moreover, lnRR and SMD diverged in heterogeneity quantification. The median *I*^2^ for the MLMA model using lnRR was 96% (mean 92%; Figure S4), while for SMD, it was 82% (mean 74%). In 31% (23/75) of cases, different interpretations of heterogeneity were possible, according to the convention (*I*^2^ = 20%, 50%, and 75% corresponding to small”, “medium”, and “large heterogeneity; Higgins et al., 2003).

For example, among 72 cases indicating large heterogeneity for the MLMA model using lnRR, 22 cases were interpreted as having moderate or small heterogeneity for SMD (Figure 3). The divergence between lnRR and SMD was also evident in the results of publication bias tests (Figure 3). While 24% (18/75) of cases were identified as having publication bias for the MLMA model using lnRR, a higher percentage, 76% (57/75), exhibited publication bias for SMD. Similar discrepancies are observed in the results from the BMLMA of lnRR and SMD (Figure S2).

The observed discordance stems from the two metrics’ different biological and statistical assumptions. We offer five insights to clarify these distinctions. First and foremost, from a biological sense, lnRR and SMD assume distinct natures of the environmental and biological effects. While lnRR expresses the mean difference at the multiplicative scale, SMD expresses the mean difference at the additive scale.

Although the multiplicative effect measured by lnRR can be transformed into the additive effect at the logarithmic scale (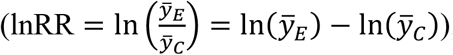 ), this variable transformation can alter the interpretation of the variable (Rönkkö et al., 2022). Secondly, lnRR has been reported to have greater statistical power than SMD (Yang, Sánchez-Tójar, et al., 2023). We found a median power of 81% (mean 63%) for lnRR and 66% (mean 58%) for SMD across our 75 meta-analysis datasets (Figure S5). Thirdly, the SMD effect size parameter assumes homoscedasticity (Equation 3; Hedges, 1981; Viechtbauer, 2007b), implying a common variance between the environmental stressor group and the control group. However, heteroscedasticity (unequal variances) was observed in 31% of cases in our database (Figure S6; Supplementary Materials).

Fourthly, SMD exhibits undesirable statistical properties, with an inherent correlation between the effect size estimator and its sampling variance estimator (Equation 5), affecting the accuracy of the overall effect estimate. Specifically, the error in estimating SMD naturally propagates into its sampling variance, or weights under the inverse-variance weighting scheme (Hamman et al., 2018). Finally, the intrinsic correlation between point estimates and sampling variances in SMD also contributed to the seemingly prevalent publication bias (as high as 76%) detected by Egger’s test (Pustejovsky et al., 2019), which assumes smaller studies (thus large sampling variances) reports larger effect sizes; SMD inherently has this correlation. Finally, a simulation study indicates substantial positive bias in existing estimators for the between-study variance for lnRR (Bakbergenuly et al., 2020), potentially inflating heterogeneity (measured as *I*^2^), as observed in our database. Additionally, the typical sampling variance in lnRR was, on average, 45 times smaller than that in SMD (Figure S7), which could also contribute to the observed large *I*^2^ observed in lnRR.

Notably, the proposed bivariate synthesis of lnRR and SMD is distinct from that of SMD and its mathematical transformation, for example, Fisher’s r-to-Zr transformed correlation coefficient (*Zr*; Nakagawa et al., 2007). This is despite the fact that lnRR, SMD, and Zr and their corresponding sampling variances can all be estimated using the same information (i.e., sample means, standard deviations, and sample sizes; Nakagawa et al., 2007). The bivariate model of SMD and *Zr* (estimated from the same data) does not include any new information because *Zr* is a direct transformation (i.e., there is a perfect correlation between SMD and *Zr* under certain assumptions of the mathematical transformation formula). Therefore, we should avoid using such a bivariate model. In contrast, as mentioned earlier, there is no one-to-one transformation between lnRR and SMD. While lnRR is a multiplicative effect, SMD is an additive effect, incorporating the information of population standard deviation.

Therefore, lnRR and SMD contribute unique information to the model, albeit they are well correlated, and, therefore, there is much new information to be gained via a bivariate model.

### 3.3 Gain in statistical and biological information

The bivariate synthesis of lnRR and SMD through the BMLMA improved the precision of the overall effect estimate for 83% (62/75) cases of lnRR and 39% (29/75) cases of SMD, compared to the MLMA of lnRR together with SMD. On average, the borrowing of strength (BoS) was 0.25 for lnRR and 0.27 for SMD (Figure 5), indicating that relying solely on one effect size (lnRR or SMD) would lead to a loss of nearly 20% and 30% of information, respectively. This incremental precision stemmed from the high correlation between lnRR and SMD effect size parameters (Copas et al., 2018), with a median correlation of 0.76 at the within-study level (Figure S8). The information gained from the bivariate synthesis of lnRR and SMD was equivalent to the benefit of discovering evidence for an environmental and biological effect from approximately 20 to 40 additional effect sizes (Figure 5).

**Figure 5.**
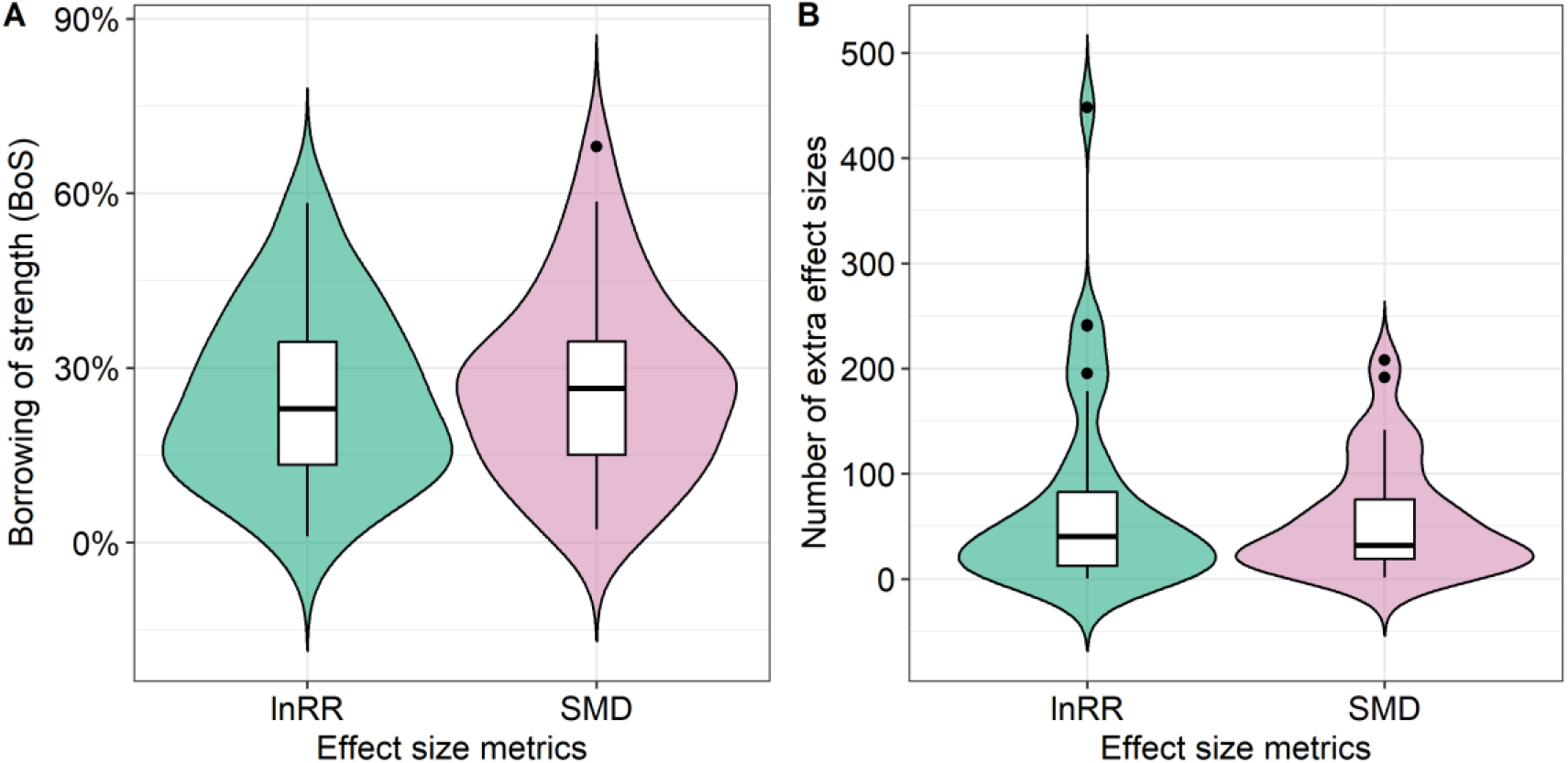
Statistical information gain resulting from the concurrent synthesis of lnRR and SMD utilizing the bivariate multilevel meta-analysis (BMLMA) model. Two information indices were used, including the borrowing of strength (BoS), representing the reduction in uncertainty or increase in precision (Panel A), and the number of additional effect sizes incorporated due to the decrease in uncertainty (Panel B).

From a biological perspective, the simultaneous use of lnRR and SMD through the MLMA model revealed that 62% of the observed variability in lnRR (46% in SMD; Figure 6) could be attributed to ‘true’ biological or methodological differences at the within-study level. Notably, our findings unraveled a compelling trend wherein nearly three-quarters of instances exhibited smaller between-study level heterogeneity, characterized by a median *I*^2^ of 23% for lnRR and 8% for SMD, compared to within-study level heterogeneity (median *I*^2^of 70% for lnRR and 61% for SMD). The practical ramifications of these observations are two-fold. Firstly, variables specific to within-study-level characteristics are more likely to predict the effect of interest than those associated with between-study-level characteristics (Viechtbauer, 2007a).

**Figure 6.**
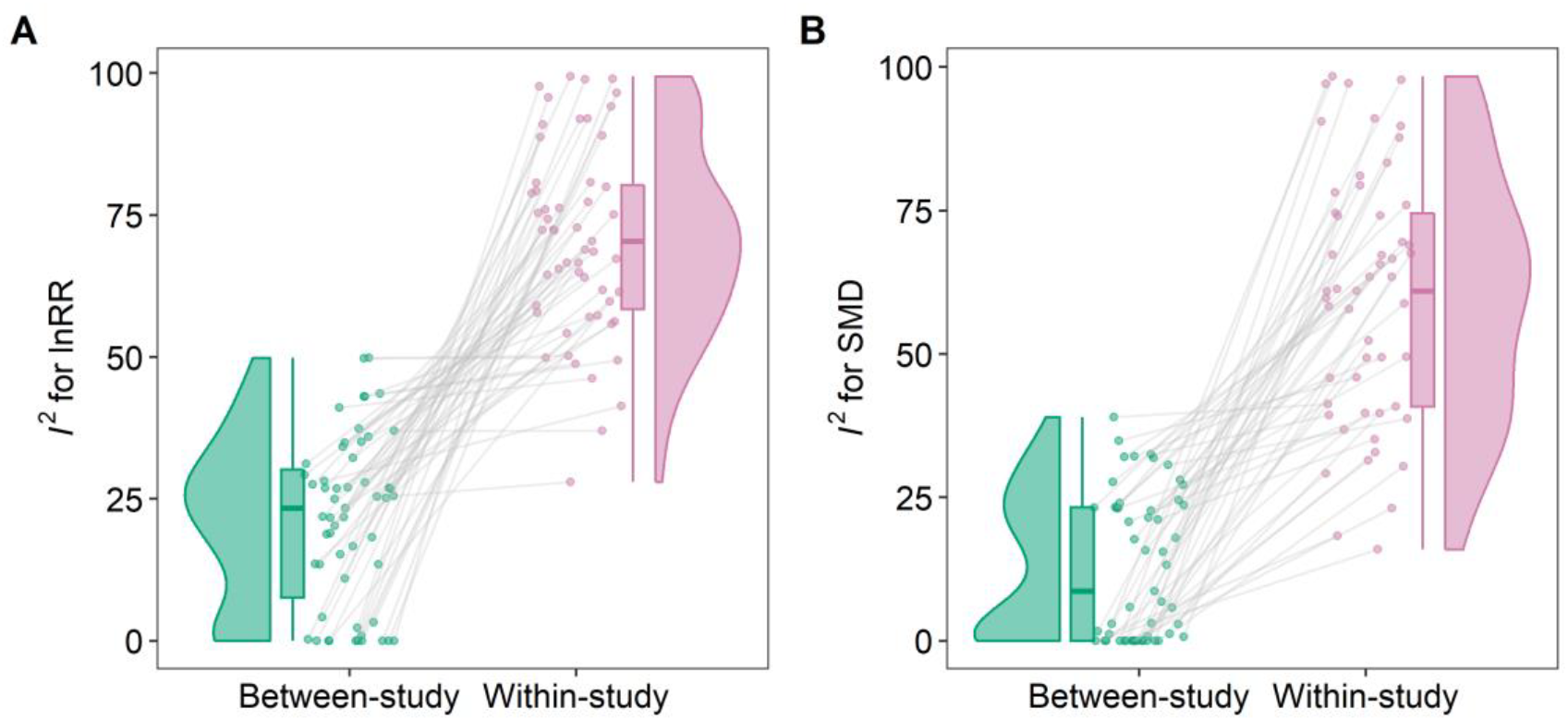
Biological information gain resulting from the multilevel meta-analysis (MLMA) of lnRR along with SMD. Heterogeneity decomposition was used to measure the gain in biological insights. The total heterogeneity surrounding the overall effect was partitioned into between- and within-study levels for lnRR (Panel A) and SMD (Panel B), facilitating the identification of predictors associated with each level.

Secondly, if within-study variations are properly controlled, generalization of research findings becomes not only achievable but also common (Yang, Noble, et al., 2023), as was evident in at least 73% of the current 75 datasets.

## 4 CONCLUSIONS

Conventionally, researchers often employ a random-effects (RE) model using only one effect size metric alone, lnRR or SMD. However, as demonstrated, this conventional practice could yield substantial erroneous conclusions and nonnegligible discrepancies in detecting and estimating the effect of interest. Occasionally, using only lnRR or SMD alone led to inconsistencies in the sign of the effect of interest. Furthermore, lnRR and SMD exhibited large discrepancies in heterogeneity interpretation and publication bias test. To address these issues, we propose the use of either a multilevel meta-analysis of lnRR and SMD together (the first advanced approach) or a bivariate multilevel meta-analysis analysis of lnRR and SMD (the second advanced approach). The two approaches can offer statistical and biological insights by leveraging the strengths of each approach.

The following two points should be noted when applying the two proposed approaches. First, the two approaches should be used in conjunction with robust variance estimation to ensure valid statistical inferences. Second, in cases where the number of included studies is small, the second approach is highly parameterized and may be prone to overparameterization. The benefits might be compromised by the increase in the number of parameters that need to be estimated (Boca et al., 2017; Yang, Macleod, et al., 2023). In such a case, we recommend using the first approach as the main method and the second as a sensitivity analysis. Finally, to enhance the accessibility of these two approaches, we have established a user-friendly website with a step-by-step implementation guide. The widespread use of our proposed method can contribute to reaching credible, reliable, and reproducible environmental and biological syntheses.

## AUTHOR CONTRIBUTIONS

**Yefeng Yang**: Conceptualization; formal analysis; methodology; visualization; writing – original draft; writing – review and editing. **Coralie Williams**: Conceptualization; methodology; visualization; writing – review and editing. **Alistair M. Senior**: Conceptualization; methodology; writing – review and editing. **Kyle Morrison**: Conceptualization; methodology; visualization; writing – review and editing.

**Lorenzo Ricolfi**: Conceptualization; methodology; visualization; writing – review and editing. **Jinming Pan**: Conceptualization; methodology; funding Acquisition; supervision; writing – review and editing. **Malgorzata Lagisz**: Conceptualization; methodology; supervision; visualization; writing – review and editing. **Shinichi Nakagawa**: Conceptualization; methodology; supervision; visualization; writing – review and editing.

## ACKNOWLEDGMENTS

YY and JP were funded by the National Natural Science Foundation of China (NO. 32102597) and the China Agriculture Research System (CARS-40). SN and ML were funded by the Australian Research Council Discovery Grant (DP210100812 & DP230101248).

## DATA AND CODE AVAILABILITY STATEMENT

The data and code that reproduce the findings of this study are openly available at https://github.com/Yefeng0920/BRMA_MD. The website that shows the step-by-step implementation of the two proposed approaches is openly available at https://yefeng0920.github.io/BRMA_tutorial_git/.

